# Density-dependent effects are the main determinants of variation in growth dynamics between closely related bacterial strains

**DOI:** 10.1101/2021.05.03.442413

**Authors:** Sabrin Hilau, Sophia Katz, Tanya Wasserman, Ruth Hershberg, Yonatan Savir

## Abstract

Although closely related genetically, bacterial strains belonging to the same species show significant variability in their growth and death dynamics. However, our understanding of the underlying processes that lead to this variability is still lacking. Here, we measured the growth and death dynamics of 11 strains of *E. coli* originating from different hosts and developed a mathematical model that captures their growth and death dynamics. Our model considers two environmental factors that determine growth dynamics: resource utilization efficiency and density-dependent growth inhibition. Here we show that both factors are required to capture the measured dynamics. Interestingly, our model results indicate that the main process that determines the major differences between the strains is the critical density at which they slow down their growth, rather than maximal growth rate or death rate. Finally, we found that bacterial growth and death dynamics can be reduced to only two dimensions and described by death rates and density-dependent growth inhibition alone.

**Importance:** Understanding the dynamics of bacterial growth has been an area of intense study. However, these dynamics have often been characterized through the narrow prism of describing growth rates, without considering parameters that may modulate these rates. Here, we generate a model that describes bacterial growth and death dynamics, incorporating two essential, growth-modulating factors: density-dependent reductions in growth rates and resource utilization efficiency. This model allows us to demonstrate that variation in the growth curves of closely related bacterial strains can be reduced to two dimensions and explained almost entirely by variation in the cellular density at which bacteria slow down their growth, combined with their death rates.

## Introduction

Bacterial growth and death dynamics, following introduction into new media, are often described through a curve composed of four distinct phases. In the first phase, bacteria prepare for growth within their new media and so experience a ‘lag’ in their ability to grow. Following this short period of lag, bacteria begin to grow in a phase often referred to as log phase. Despite its name, bacteria only initially grow exponentially and then slow to sub-exponential growth. Following growth, once bacteria reach their maximal yield, they enter stationary phase, a phase in which growth and death rates are equal, leading to maintenance of consistent viability with time. The stationary phase cannot be maintained indefinitely, and so bacteria enter the death phase, during which they undergo an exponential loss of viability^1–5^.

Growth and death dynamics are known to vary significantly between different bacterial species. This variability is reflected in several parameters. The most well studied so far being variation in the maximal doubling time of different bacterial species. For example, *Escherichia coli* undergoes cell replication approximately every 20 minutes, *Pseudomonas aeruginosa*, replicates about every 30 minutes, whereas *Syntrophobacter fumaroxidans* can replicate only once every 140 hours^6,7^. Closely related strains of the same species can also display phenotypic diversity in various properties, such as sensitivity to various stresses^8^, resistance to antibiotics^9^ and cell size and shape^10^. Yet, variation in growth dynamics between closely related bacterial strains has not been well characterized.

Adaptation of complex dynamics often involves tradeoffs between various phenotypic parameters that together result in those dynamics^11,12^. In the context of growth dynamics, the tradeoff between maximal yield and maximal growth rate is the most well investigated^13,14^. It was suggested that the relationship between maximal growth rates and yields follows a “bell curve”, divided into two limbs. The first limb encompasses the lower range of maximal growth rates up to a certain threshold. In this range, maximal growth rate and yield are positively correlated. Within this range, resources are sufficient to support growth and as strains become more efficient in utilizing resources they can grow faster and reach higher yields. The second limb of the curve encompasses strains with higher maximal growth rates. In this range, the maximal growth rate and yield become negatively correlated, because cells do not have sufficient capacity to support both their rapid growth and the metabolic requirements for maximizing yield^13^.

In contrast to parameters related to growth, parameters related to death are often neglected^15^. Death rates of various bacterial strains and species are very rarely characterized, at least partially due to technical difficulties resulting from the widespread use of optical density measures of cell growth, that cannot capture reductions in cell numbers for most species. As a result, tradeoffs between growth and death parameters are ill characterized.

Mathematical models are widely used for understanding and predicting bacterial behavior under different environmental conditions (such as in environments that vary in their pH or temperature)^16,17^. Bacterial growth kinetics were previously described by a large variety of mathematical models^17,18^. While most of these models describe the first three phases of the bacterial growth curve, only a few models include the death phase and present it separately from growth^19,20^.

Bacterial growth is often limited by resource availability. The response of bacterial growth to resource limitation is often captured in models of growth by the Monod equation that describes the relationship between resource availability and growth rates^21^. At the same time, bacterial growth can also be limited by intrinsic mechanisms that reduce growth, once populations reach certain densities. The signal-response mechanism that enables bacteria to synchronize growth at the population level and alter their growth in response to increases in cellular density is known as quorum sensing (QS)^22–25^. QS is based on the production and secretion of molecules, called autoinducers, to the medium. These autoinducers accumulate as a function of population density, reaching a minimal threshold concentration detected by the bacteria. Once this concentration is reached, bacteria alter their gene expression, leading to changes in growth ^26^. Growth rate is one of the parameters that declines drastically in response to increased density^27^. While density-dependent effects substantially affect growth dynamics, they are not usually parameterized in mathematical models of growth dynamics.

Here, we develop a novel mathematical model of bacterial growth dynamics that parameterizes both resource limitation and density-dependent effects and that considers all four stages of the bacterial growth curve. We apply this model onto data we collected of nine natural *E. coli* isolates and two *E. coli* lab strains and demonstrate that the model captures their growth dynamics. We show that bacterial population dynamics cannot be well described without including parameters describing *both* the density-dependent and resource-limitation-dependent effects on growth. Applying our model on data shows that variation in maximal yield between strains is determined mainly by the cellular density at which growth rate is reduced, rather than by the maximal growth and death rates. We also show that across *E. coli* strains, strains that slow down growth at higher cellular densities tend to utilize their resources more efficiently. Finally, we show that because maximal doubling time is fairly consistent across strains and due to the correlation between the density dependent effect and resource utilization efficiency, the growth dynamics of any specific strain can be well-characterized using only its death rate and density-dependent effect.

## Results

### Different *E. coli* strains show significant variability in their growth and death dynamics

Various *E. coli* strains populate different environments that vary in their evolutionary pressures and types of stress. As a result, the strains adapt to their environment by optimizing their growth and death dynamics according to environmental demands. To obtain growth data for 11 *E. coli* strains originated from different environments, we performed both optical density (OD) and colony forming units (CFU) measurements. OD measurements are a proxy for the total number of bacteria (live and dead) present within a sample. OD measurements enable fine-grained estimation of growth rates, but do not enable one to quantify death rates, as dead cells and their debris, continue to be measured. To measure CFU, cells are plated onto agar plates and the resulting colonies are counted. CFU measurements allow the quantification of death rates, as they do not account for dead cells. We quantified OD for each of the 11 strains by carrying outgrowth experiments within a plate reader for the first 16 hours of growth, measuring OD at 10-minute intervals (Methods section). For CFU measurements we grew bacteria for 56 hours, sampling and quantifying CFUs at 11 time points (0,1,2,3,8,16,24,32,40,48,56 hr.) (Fig. 1A, Table S1, Methods section). Figure 1A depicts the growth curves extracted, based on CFU quantification, for two of the 11 considered strains and exemplifies the clear variation observed between strains (Fig. 1A).

**Figure 1:**
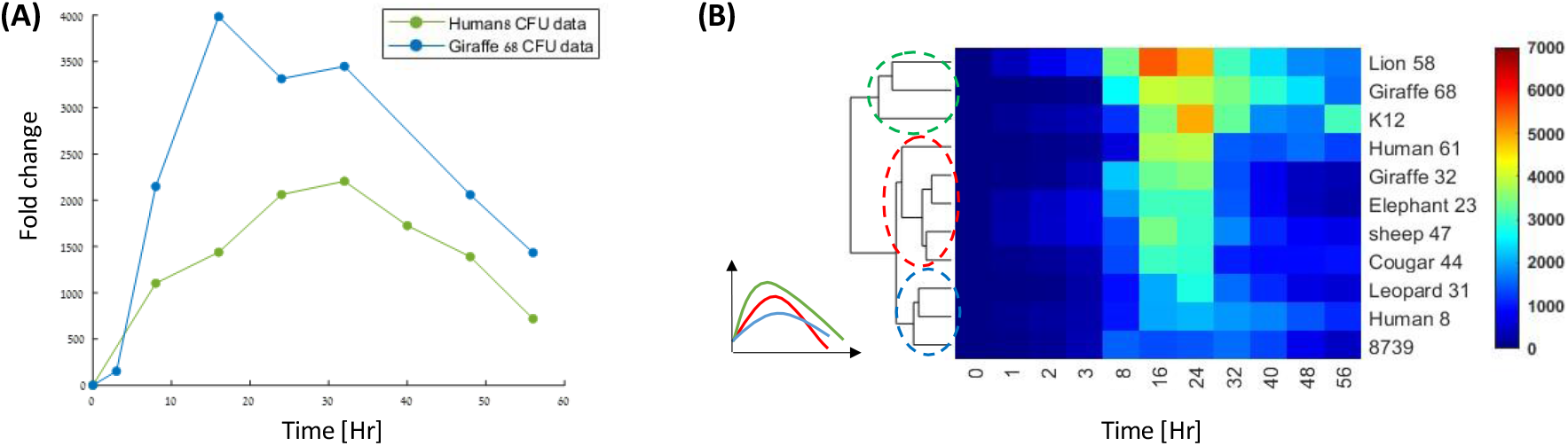
*E. coli* strains show significant variability in their growth and death dynamics. **(A)** Two representative growth curves of two samples from different natural strains. The dots represent the bacteria live count in a specific time point normalized by the starting point. **(B)** Growth dynamics hierarchical clustering of 11 *E. Coli* strains isolated from various environments reveals three groups that exhibit different dynamics patterns. The analyzed time series vectors are the median trajectories of three to seven samples for each time point.

To validate and estimate the significance of the variation observed between the growth curves of each pair of the 11 strains, we treated each curve as a time series and performed hierarchical clustering analysis on their median time trajectory (Fig. 1B). The clustering reveals three main groups of strain growth curves with different dynamical properties. The first group (green, Fig. 1B) is characterized by high maximal yield and slow death, the second group (blue, Fig. 1B) is characterized by low maximal yield and slow death, and the third group (red, Fig. 1B) has intermediate maximal yield values and can be further divided into two groups that are differed by their death rate.

To characterize the relationship between the observed maximal growth rate, death rate, and maximal yield, we derived the maximal growth rate and the death rate for each strain, from the growth curve data (see Supplementary Information, sections II, III). A surprisingly low variation in maximal growth rates was observed between strains (Figure 2A). Only two strains (Human 8 and the K12 lab strain) varied from all remaining nine strains in their median maximal growth rates. In contrast to maximal growth rates, maximal yields and death rates varied more substantially between strains. As a result, we observed no correlation between maximal growth rate and either death rates (Figure 2B), or maximal yields (Figure 2C). Finally, death rates also do not correlate significantly with maximal yield (Figure 2D).

**Figure 2:**
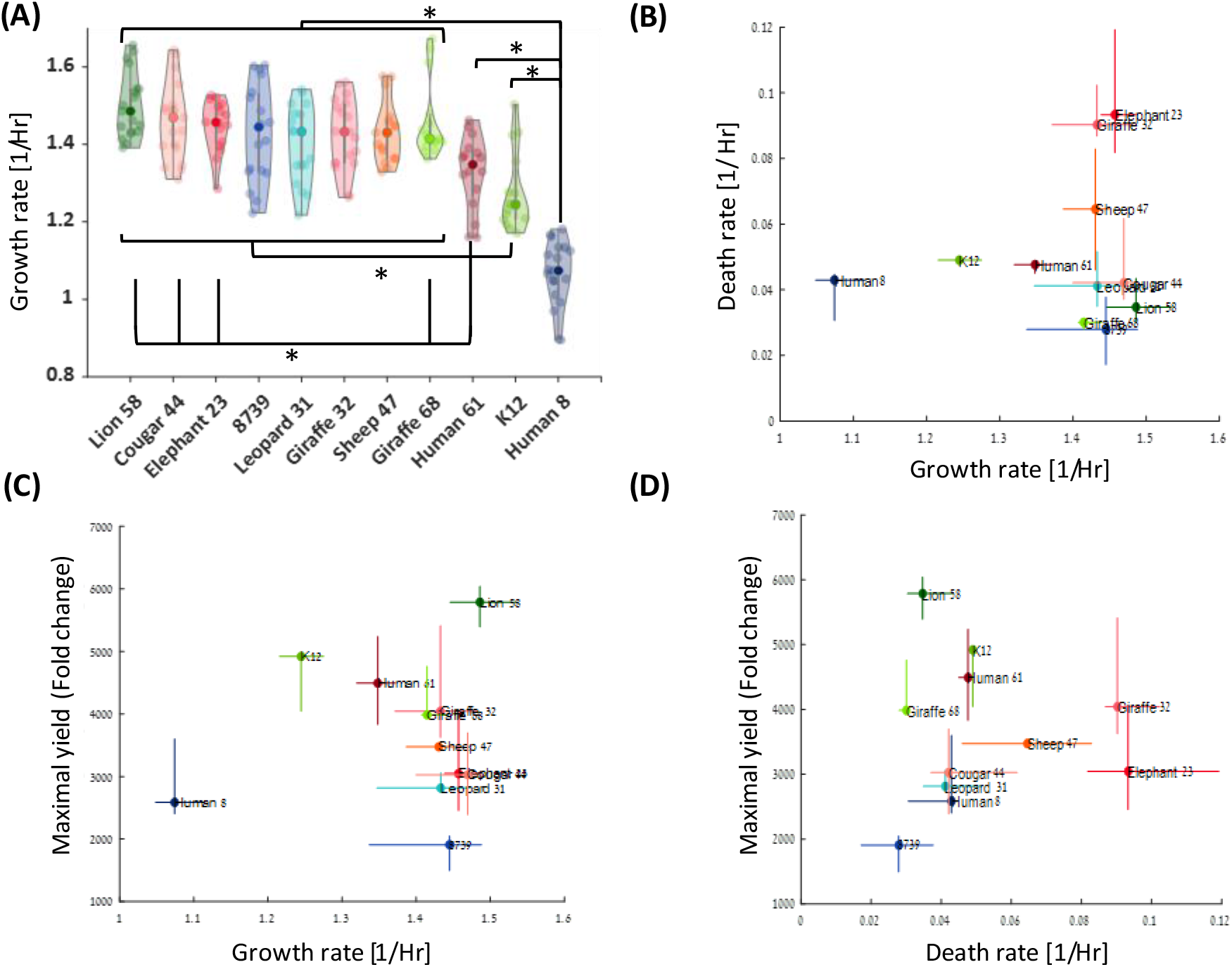
Maximal growth rate and death rate do not account for strain variability. **(A)** Violin plot of the maximal growth rate distribution. The bold dots represent the median maximal growth rate. The other dots represent the different samples. The asterisk denotes distributions that are different according to the Kolmogorov-Smirnoff test with a p-value<0.05. **(B)** Maximal growth rate vs. death rate. The colored dots represent the median rates. The error bars are the 34 and 66 percentiles. Maximal growth rate and death rate have no significant correlation between them. There is also no significant correlation between **(C)** Maximal yield and growth rate and **(D)** Maximal yield and death rate.

### A mathematical model that takes only resource limitation into account cannot capture the observed variation in growth and death dynamics

The Monod model, which takes into account resource limitation as a growth rate modulator is commonly used for modeling bacterial growth^23,28^. Following Monod, we developed a mathematical model that integrates growth, death and resource limitation (Fig. 3A). The model is composed of two coupled ordinary differential equations (ODE) for the dynamics of bacteria number, *N*, and the resource, *r*,

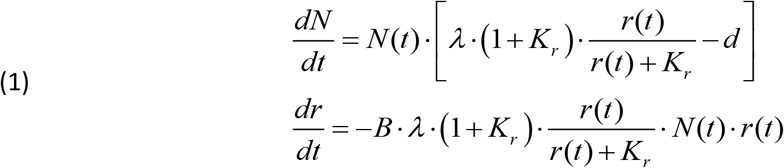

where λ is the maximal growth rate, *d* is the death rate, *B* is the number of resource units the bacterium needs to divide once, and *K_r_* is the resource amount at which the growth rate is half the maximal growth rate. We have normalized the resource value at time zero to be one, *r*(0) = 1.

**Figure 3:**
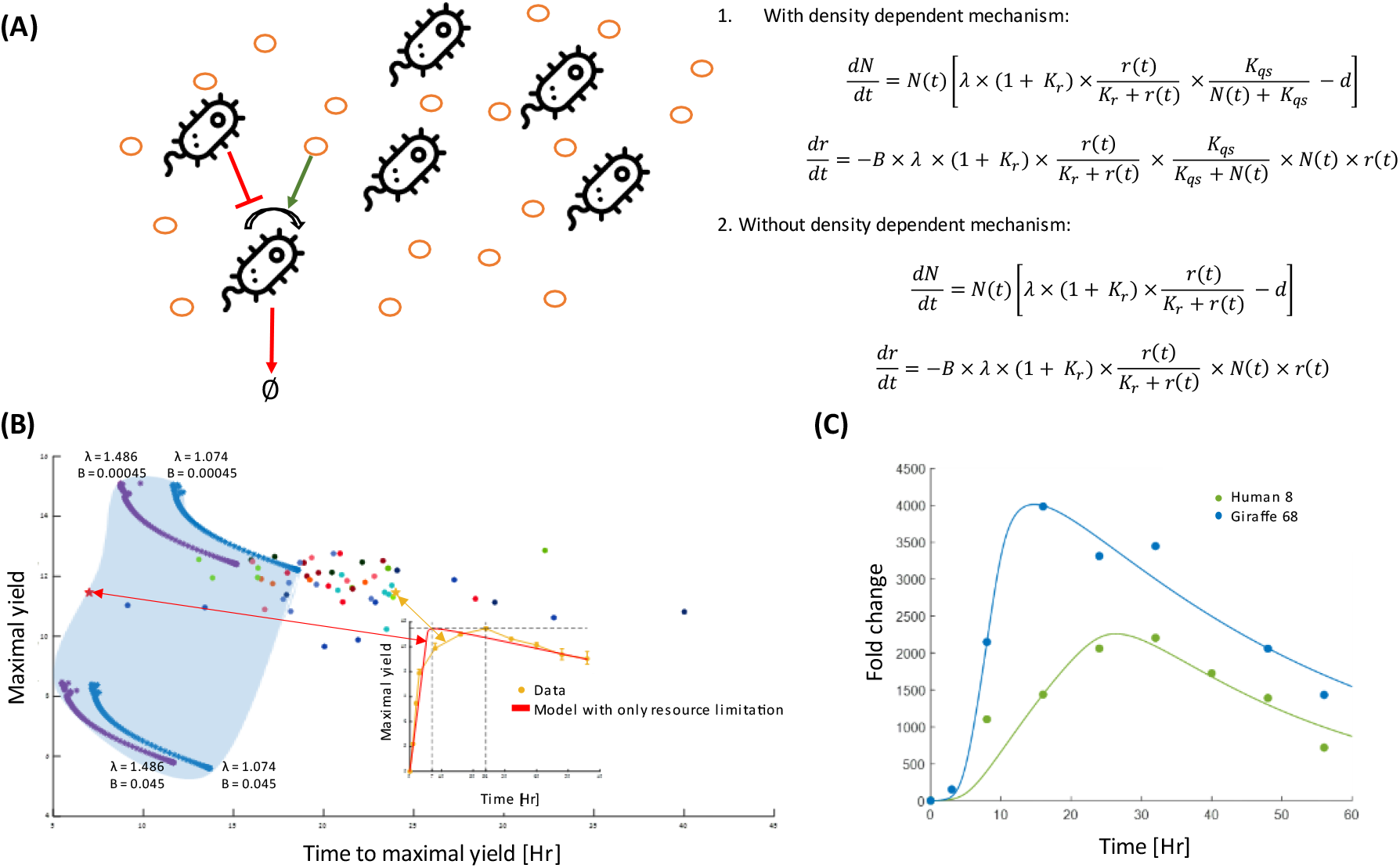
Density-dependent limitation and resource limitation are both necessary to capture the entire growth dynamics. **(A)** Growth and death dynamics regulation is composed out of two environmental factors and described by two coupled ODEs. The orange circles denote the resource. The resource is needed for bacterial growth (green arrow), and bacterial density negatively affects growth (red arrow). The first two coupled ODEs describe these dynamics and include two growth rate modulators: resource amount and bacterial density. The second pair of ODEs describe bacterial growth dynamics considering only resource limitation. **(B)** Quantifying the models’ limits. The cyan area is the possible values of maximal yield and time to maximal yield from the model that takes only resource limitation into account (model (2)). These values are estimated for extreme values of *λ* and *B* and a wide range of *Kr* (10^−6^-10^2^) (blue and purple asterisks). The model with only resource limitation cannot capture the experimental data (full circles). The inset illustrates an example for an experimental measurement that is outside the resource limitation feasible space (yellow star). The red curve is the recourse limitation model that has the same maximal yield as the data. Note that for a model that accounts only for resource limitation, the time to maximal yield, for a given maximal yield, is much shorter than observed. **(C)** A representative fit of the model with both resource limitation and density-dependent effect on the data. The presented samples are the two samples that were shown in Fig 1A.

We find that this model cannot capture the entire observed growth and death dynamics (Fig. 3B). One of the main limitations of this model is that it predicts that a population will reach maximal yield faster than it actually does. Figure 3B illustrates the phase space of the relationship between the time to reach maximal yield and the value of the maximal yield, for a broad set of parameters. Most of the experimental observations fall outside of this regime. Thus, this model cannot capture the variation in the observed dynamics between strains, not because of poor parameter fitting, but rather due to the inherent structure of the model.

### Density-dependent effects determine maximal yield variability

Since a model that considers only resource limitation cannot capture our observed growth dynamics, we added to the model another term that modulates growth in a density-dependent manner,

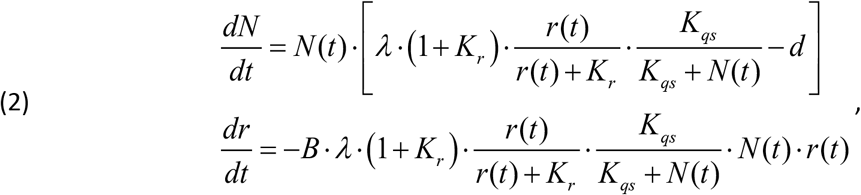

where *K_qs_* is the bacterial density at which the density-dependent term reaches half its maximal value and reduces the growth rate dramatically. Fitting the data with the modified model, we showed that the density-dependent term enables us to capture the data more accurately (Fig. 3B, 3C). The interplay between the resource and density-dependent effects on growth is such that growth rate decline due to the density effect is dominant earlier in the dynamics than resource limitation (Supplementary Information section IV, Fig. S1). Since the death rate is negligible compared to the maximal growth rate, the maximal yield, and the time it takes to reach it, are not strongly influenced by death rates (Supplementary Information section IV, Fig. S1).

While growth rate and death rate are not correlated with maximal yield (Fig. 2C, 2D) the bacterial density at which the growth rate is reduced drastically, *K_qs_*, and the number of divisions bacteria can undergo on a certain amount of resource units, 1/*B*, are significantly correlated with the maximal yield and with each other (Fig. 4). In other words, strains that achieve higher maximal yields, use their resources more efficiently and delay their density-dependent slowdown, relative to strains that reach lower maximal yields. Interestingly, the groups that were identified using the non-parametric clustering keep their structure in the phase space of these three kinetic parameters of the model (Fig. 4). The blue cluster is composed out of strains that grow most slowly and have the lowest maximal yields (Fig. 1B). Strains belonging to this cluster also appear to be the most inefficient in terms of resource utilization and slow their growth rate after reaching the lowest cellular density. In contrast, strains belonging to the green cluster are characterized by the highest maximal yields, are very efficient in their resource utilization and delay growth slowdown when they reach much higher densities. Strains belonging to the red cluster have intermediate maximal yields. Fitting with this, their resource utilization efficiency and the density at which they reduce growth rates are also intermediate (Figures 1B, 4A).

**Figure 4:**
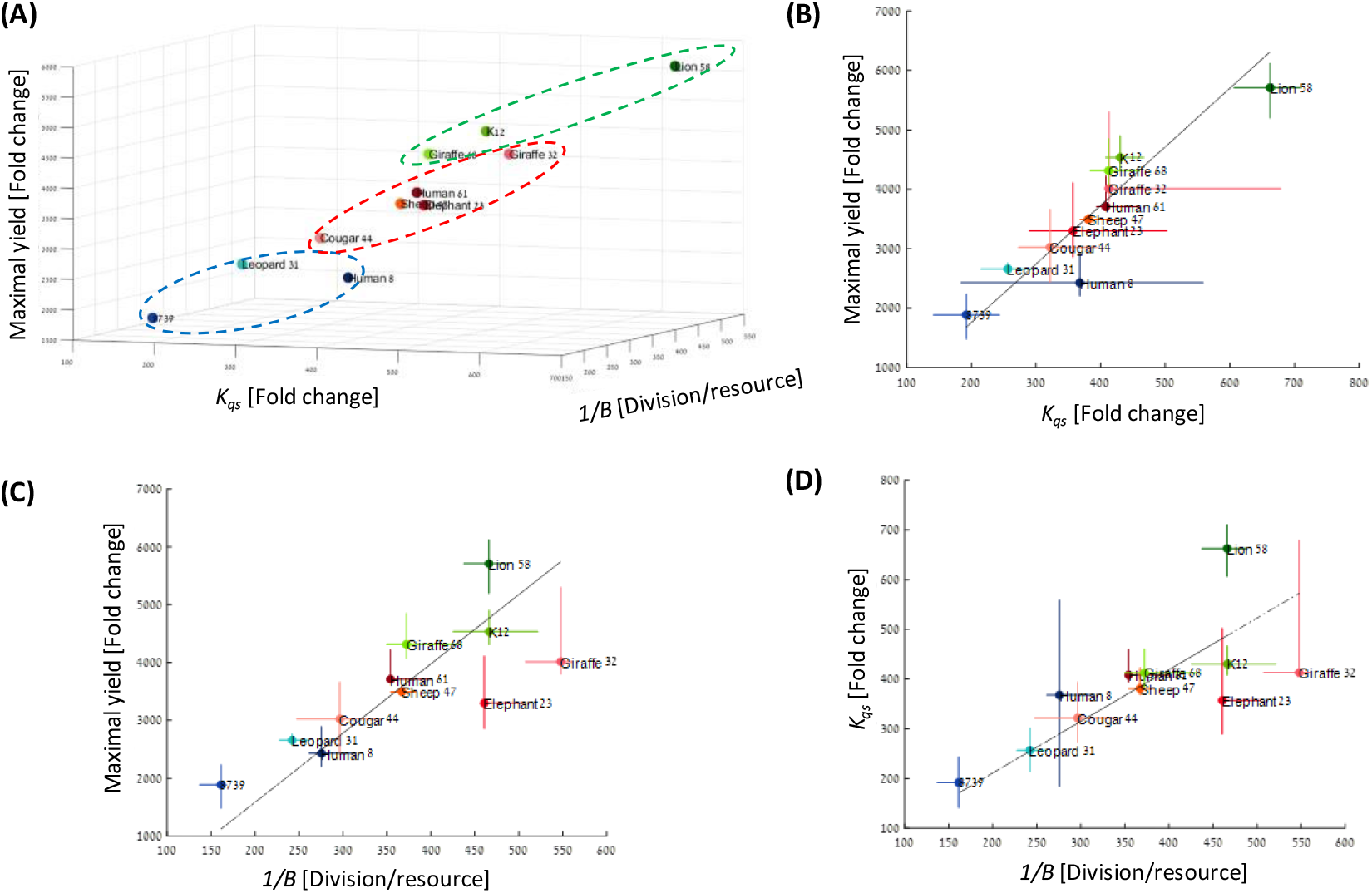
Maximal yield is correlated with density dependent effect. **(A)** Maximal yield vs. *K*_qs_ and 1/*B*. The dots represent the median of each strain. **(B)** Maximal yield vs. *K*_qs_. The gray line is the best linear fit (*ρ*= 0.94; P-value<0.001, where *ρ* is spearman correlation). **(C)** Maximal yield vs. 1/B (resource utilization efficiency). (*ρ* = 0.85; P-value= 0.0016). **(D)** 1/B vs. *K*_qs_. (*ρ* = 0.84; P-value= 0.0021).

### Strains reside within a two-dimensional phenotypic space

While the model contains five parameters, the significant correlation between the density-dependent effect (*K*_qs_) on the maximal growth rate (*λ*) and resource utilization efficiency (*1/B*) suggests that the effective dimension of the phenotypic space is lower. To examine the number of parameters necessary to describe the bacterial growth and death dynamics, we performed principal component analysis (PCA). Within our PCA, each strain was characterized by a vector composed of the five model parameters (maximal growth rate (λ), death rate (*d)*, the density-dependent effect (*K*_qs_), resource utilization efficiency (1/*B*), and the resource concentration at which growth rates reach half of λ (*K*_r_). 72.5% of the variability is explained by the first two principal axes, while >90% are explained by the first three. Figure 5 illustrates the phenotypic distribution in the death rate, efficiency and *K*_qs_ space. The data mostly reside within a two-dimensional space plane spanned by the first and second PCA axes, A1 and A2 (Fig. 5A, 5B). The structure of the data within this plane is intriguing as it reveals a phenotypic distribution that is close to a triangle. (Fig. 5C, 5D). This type of dimensional reduction may indicate an adaptation process where the nodes of the triangles represent an archetype combining all three traits at different proportions. Each archetype is adapted to specific type of environment^11,12^ (Fig. 5C, 5D). Interestingly, each one of the different groups identified earlier, populates different nodes of the triangle. The strains on the blue edge experience a density-dependent slowdown at low densities and are inefficient in their utilization of resources. However, their death rates are the lowest. Strains on the red edge die very rapidly but are very efficient and insensitive to bacterial density. The green edge strains slow down their growth at the highest densities and are intermediate in respect to their efficiency and death rates. To validate the PCA results and demonstrate that the strains can be separated by two parameters only (*d*, *K_qs_*, or 1/B), we evaluated the parameters that separate most significantly between the strains in the edges. To do so, we calculated the distance in the 5D phase space between the strains in the triangle nodes (Fig. 5D). We observed that the edges are separated by two to three parameters, which are a combination of *d*, *K_qs_*, or 1/B.

**Figure 5:**
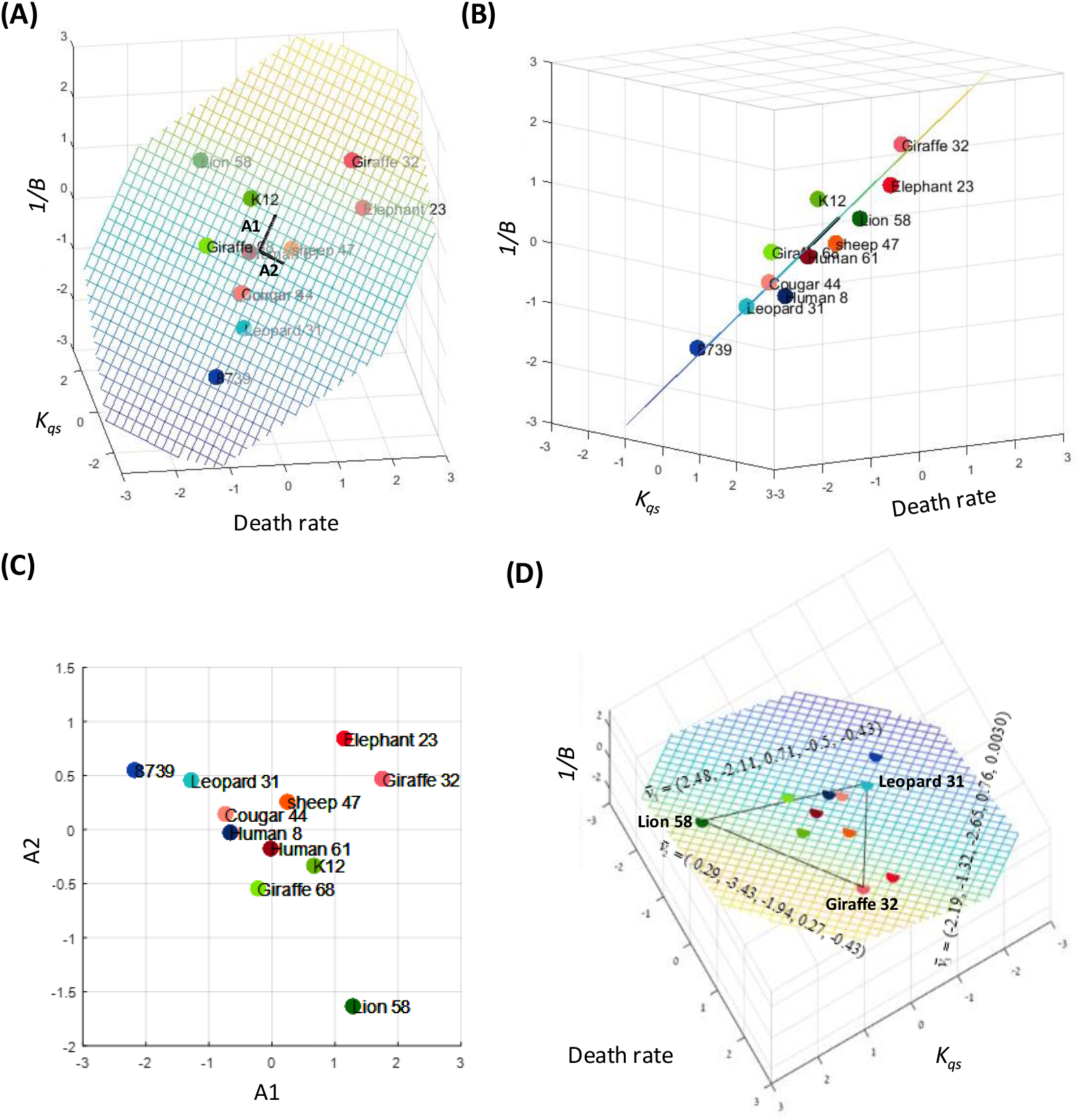
Phenotypic diversity is confined to a low dimension. **(A)** Each dot represents a strain in the three-dimensional space of *d*, *K_qs_*, and 1/*B*. The black vectors are the projection of the 5D first and second PCA axes, A1 and A2, respectively, to this space. The dashed plane is the 2D plane spanned by A1 and A2. The parameters reside mostly in this plane. **(B)** Projection of the data along with A1. The deviation of the data plots from the phenotypic plane is small compared to its spread within the plane. **(C)** The data projected onto the plane spanned by A1 and A2. **(D)** The projected data in the phenotypic plane and the phenotypic distance between representative strain on the triangle’s edges (parameters order: *d, K_qs_, 1/B, K_r_, λ*).

## Discussion

To enable us to compare the entire growth curve of different bacterial strains and to extract the parameters that most strongly influence variation between strains, we generated a mathematical model that fits experimentally generated growth curves. To allow us to fit the data, we first observed that growth rate must be modulated not only by resource limitation, but also by bacterial density, that was revealed to be the dominant factor influencing growth rate earlier in the dynamics. This likely results from resources being highly abundant and thus not limiting growth, during the initial stages of growth. However, once resources are depleted, growth rates become limited by resource availability and the resulting fits follow the Monod equation, we used in our model^29^.

Our model was used to extract three parameters in addition to more directly estimated maximal growth rate and death rates: *K_qs_* (the bacterial density in which they slow down their growth significantly), *1/B* (the resource utilization efficiency) and *K_r_* (the amount of resource at which growth rates reach half the maximal growth rate). Comparing these parameters, we found that *K_qs_* and 1/*B* are correlated with the maximal yield and with each other significantly. At the same time, maximal growth rates and death rates did not correlate with yield, and indeed maximal growth rates appeared to be mostly constant across strains. These results demonstrate that the extent to which strains are able to produce cells within the same type of rich media, depends not on their ability to grow or die faster or slower. Rather, it depends most strongly on the cellular density in which it reduces its growth and on the efficiency with which it utilizes its resources. Intriguingly, both these parameters are strongly correlated, meaning that strains that reduce growth at higher densities, also tend to be more efficient in their resource utilization.

Density dependent regulation of growth has been extensively studied^30,31^. The best characterized mechanism for density dependent behavior regulation is Quorum Sensing (QS). Bacteria that use QS to regulate their population density produce, secrete and sense at least one type of QS molecules. The QS molecules increase as a function of bacterial density to certain concentration threshold in which a behavioral change occurs^32^. In our case, the behavioral change would be growth rate decline as a means to avoid population overflow in a space and resource-limited growth conditions. Strains that slow down their growth when their populations reach higher densities can be said to be less sensitive to QS signals. According to our results, the same strains also tend to utilize growth resources more efficiently and, as a result, reach higher maximal yields. Combined, this may suggest a potential cost for QS. Strains that start to execute QS at lower bacterial densities may need to utilize more of their resources to generate more of the signaling molecules involved in QS, leading to their lower efficiency and lower maximal yield. Fitting with this idea, it was suggested that due to metabolic tradeoff between bacterial growth and signal production and the accumulation of toxic products, signal production leads to a two-fold metabolic burden under resource limitation^33^. Inversely, it is also possible that bacteria that use their resource inefficiently for other, genetic or environmental reasons, may adopt high sensitivity to population density, in order to better regulate their growth and protect themselves from extinction.

Our PCA results show that the phenotypic diversity in growth and death traits resides within a two-dimensional subspace of the five-dimensional space spanned by the model parameters. Specifically, when plotting the data within a space spanned by the density-dependent effect (*K_qs_*), resource utilization efficiency (*1/B*, which is strongly correlated with *K_qs_*), and death rate (*d*), the data resides mostly within the plane spanned by the projection of the first and second PCA axes. Moreover, the phenotypic distribution within this plane takes the shape of a triangle. According to the Pareto front concept, this type of structure was suggested to reflect the result of adaption under constraints. Each triangle node represents a desired bacterial archetype composed of values of quantitated traits. Gaining optimality of one trait is at the expense of other traits’ optimality, generating a tradeoff between them. Therefore, archetypes in the nodes are characterized by trait combinations that are optimally adapted to a specific type of environment, although not all traits necessarily hold their optimal value^11,12^.

Remarkably, the strains on the nodes of this triangle correspond with the clusters that were identified in our time-series based clustering, similar to the clusters revealed in the maximal yield-*K_qs_*-*1/B* phase space (Fig. 4A, Fig. 5, Fig. 6). Members of the first archetype (the blue cluster) can be seen as “inefficient growers / strong communicators”. Their density-dependent growth slowdown occurs at low bacterial concentration (meaning they likely respond strongly to QS), and their efficiency is low. Death rates within this cluster are low. Members of the second archetype (the red cluster), are highly efficient, but die rapidly. In the third archetype (the green cluster), density-dependent growth slowdowns at much higher cellular densities, compared to the other archetypes. Green cluster members are also highly efficient, albeit a little less so than members of the red cluster. It can thus be said that the green archetype members are inverse to the blue archetype, in that they are “efficient growers / poor communicators”. The archetypes we found here emphasize the tradeoffs between the three traits and demonstrate that bacteria adapt to their environment by evolving different combinations of these trait values.

**Figure 6:**
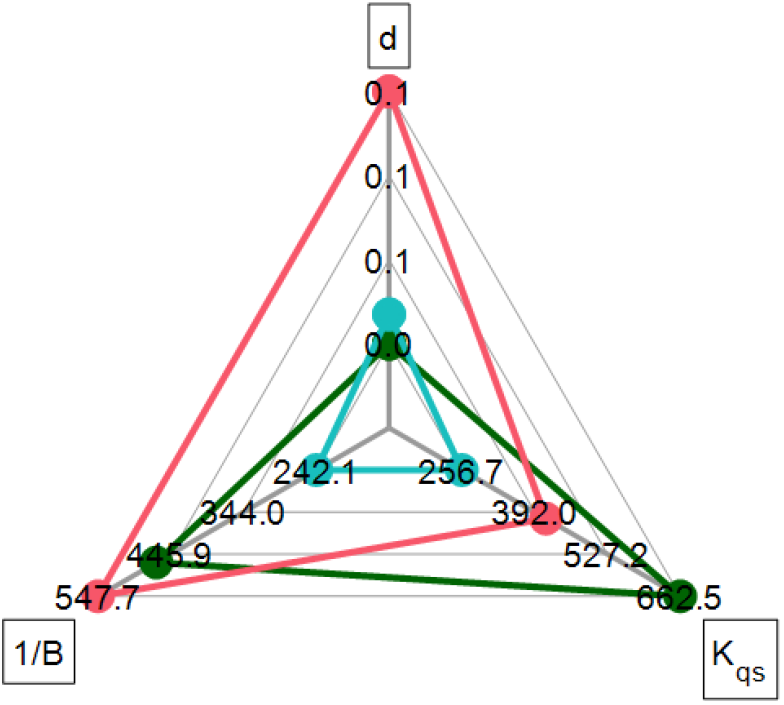
Spider plot for representative strains. Giraffe 32 (red), Lion 58 (green) Leopard 31 (cyan).

Our results emphasize the role of density-dependent growth control and its relationship with the resource allocation and maximal yield of *E. coli*. The phenotypic distribution suggests adaptation under constraints according to which the main modulators of the *E.coli* fitness are the tradeoffs between the allocation of resource to growth, its degree of interaction with its surrounding bacteria and death rates.

## Methods

### CFU measurements

Frozen samples were streaked on Luria-Broth (LB) agar Petri dishes and were incubated in 37°C overnight. Colonies were inserted to 4 ml LB in 15 ml test tube and were grown in 37°C with shaking at 225 rpm for two hours to mid-exponential stage. Samples were diluted to a concentration of 10^6^ per 2ml at a 12-well plate. Only eight wells from the plate were used for the experiment. The strains were cultured for 56 hours, and samples were taken at constant time points (at time zero, 3 hours, 8 hours, and every 8 hours). At each time point, samples were streaked by the robot on Petri dishes containing LB agar. The dishes were incubated at 37°C overnight, followed by a colony count. From each strain, five growth curves were created.

### OD measurements

Frozen samples were streaked on Luria- Broth (LB) agar Petri dishes and were incubated in 37°C overnight. Colonies were inserted to 4 ml LB in 15 ml test tube and were grown in 37°C with shaking at 225 rpm for two hours to mid-exponential stage. The test tubes were incubated in 37°C with shaking in 225 rpm for two hours to mid-exponential stage. Samples were washed twice and were diluted to 200 μl OD of 0.05 in 96-wells plate. The plate was incubated in a plate reader for 16 hours in 37°C with orbital shaking. OD was measured every 10 minutes at 595 nm. The experiment was conducted five times for each strain. Each experiment includes 3-4 samples for each strain. The OD measurements were corrected according to the plate reader calibration curve (Supplementary Information section II).

### Model fitting and model parameters evaluation

The model equations were solved numerically using “ode45” Matlab ODE solver. The simulation results were fitted to the CFU data using Matlab.

