## Supplemental information for "Density-dependent effects are the main determinants of variation in growth dynamics between closely related bacterial strains"

### Supplementary Information

#### I. Strains

In this study, we used 11 strains of *E.Coli* (Table S1). Nine of them are natural isolates obtained, as described in (1). We annotated the strains in this work according to their name and their number in the original reference list. The remaining two strains are laboratory strains.

| Strain name | Host | Number in reference list | Origin |
|---|---|---|---|
| **RM77C** | Human (Female) | 8 | Iowa |
| **RM183E** | Elephant | 23 | Washington (zoo) |
| **RM12** | Leopard | 31 | Washington (zoo) |
| **RM28** | Giraffe | 32 | Washington (zoo) |
| **RM1891** | Cougar | 44 | Washington (zoo) |
| **RM211C** | Sheep | 47 | New Guinea |
| **RM185S** | Lion | 58 | Washington (zoo) |
| **FN23** | Human (Female) | 61 | Sweden |
| **RM224H** | Giraffe | 68 | Washington (zoo) |
| **K12 MG 1655** | Laboratory strain | - | - |
| **8739** | Laboratory strain | - | - |

**Table S1:** Strains table.

#### II. Growth rate evaluation

Frozen samples were streaked on Luria- Broth (LB) agar Petri dishes, and were incubated in 37$^o$C overnight. Colonies were inserted to 4 ml LB in 15 ml test tube and were grown in 37$^o$C with shaking at 225 rpm for two hours to mid-exponential stage. The test tubes were incubated in 37$^o$C with shaking in 225 rpm for two hours to mid-exponential stage. Samples were washed twice and were diluted to 200 μl OD of 0.05 in 96-wells plate. The plate was incubated in a plate reader for 16 hours in 37$^o$C with orbital shaking. OD was measured every 10 minutes a wavelength of 595 nm. The experiment was conducted five times for each strain. Each experiment includes 3-4 samples for each strain.

To account for the nonlinearity of the OD measurements, the OD measurements were corrected according to the plate reader calibration curve. Using a plate with bacteria serial dilution, OD vs. density were measured. The resulted curve was fitted to a Michaelian function and was used to calibrate OD values,

(1) $$OD - Blank = \frac{\alpha \times C}{C + K}$$

where, α = 0.927 and K = 1.174.

The first two hours of the growth curve for each sample were fitted with an exponential growth equation,

(2) $$c(t) = c(0)e^{\lambda t}.$$

The λ values and their confidence levels were calculated from the fit (Fig. 2 in the main text).

#### III. Death rate evaluation

The CFU data were fitted to exponential decay,

(3) $$N(t) = N(t_{\max})e^{-d(t-t_{\max})},$$

where $N(t_{\max})$ is the maximal CFU, and $d$ is the death rate. The $d$ values and their confidence levels were calculated from the fit (Fig. 2 in the manuscript).

#### IV. Temporal behavior of the different growth terms

Our model contains two terms that modulate the growth (Eq. (2) in the manuscript). One growth term is modulated by resource availability, $M_r = (1+K_r)\cdot\frac{r(t)}{r(t)+K_r}$, and the other growth term is modulated by the bacteria density, $M_d = \frac{K_{qs}}{K_{qs}+N(t)}$. Figure S1 shows the typical behavior of these terms as a function of time. The main effect that determines the overall growth rate is the density effect. The decline of the resources dependent term is

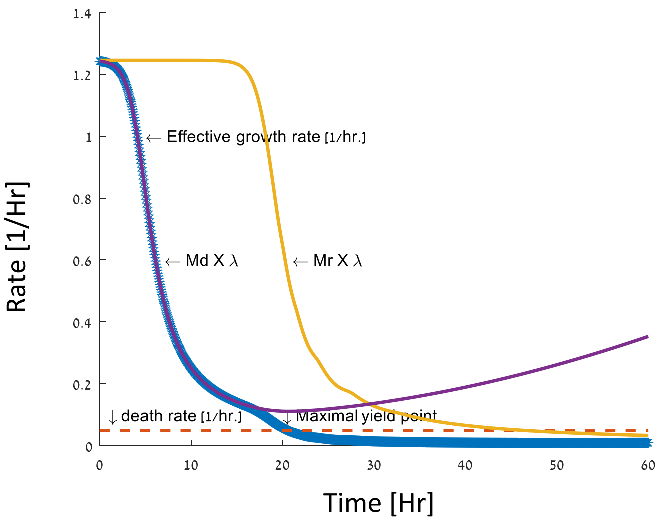


**Figure S1: The growth rate terms and death rate as a function of time.** A typical realization of the change in growth terms with time. The growth terms that depend on density (purple) and on resource decline (yellow) with time. The density-dependent term effect first and responsible for the initial, slow, decline in the overall growth rate. The resource limitation term declines fast, as expected. Both growth terms affect the growth in the range where the death term is negligible.

sharp, as expected. Both effects shape the growth before the death rate becomes non-negligible.
